# Newly synthesized RNA Sequencing Characterizes Transcription Dynamics in Three Pluripotent States

**DOI:** 10.1101/2021.06.11.448016

**Authors:** Rui Shao, Banushree Kumar, Katja Lidschreiber, Michael Lidschreiber, Patrick Cramer, Simon J. Elsässer

## Abstract

Unique transcriptomes define naïve, primed and paused pluripotent states in mouse embryonic stem cells. Here we perform transient transcriptome sequencing (TT-seq) to *de novo* define and quantify coding and non-coding transcription units (TUs) in different pluripotent states. We observe a global reduction of RNA synthesis, total RNA amount and turnover rates in ground state naïve cells (2i) and paused pluripotency (mTORi). We demonstrate that elongation velocity can be reliably estimated from TT-seq nascent RNA and RNA polymerase II occupancy and observe a transcriptome-wide attenuation of elongation velocity in the two inhibitor-induced states. We also discover a relationship between elongation velocity and termination read-through distance. Our analysis suggests that steady-state transcriptomes in mouse ES cells are controlled predominantly on the level of RNA synthesis, and that signaling pathways governing different pluripotent states immediately control key parameters of transcription.

## INTRODUCTION

Pluripotency in the pre-implantation embryo is of transient nature *in vivo*, but embryonic stem cells can be cultured long-term in stable and interconvertible pluripotent states *in vitro*: in serum/leukemia inhibitory factor (LIF) media (SL, serum-primed state), serum-free media containing LIF, Mek1/2 and GSK3*β* inhibitors (2i, naïve/ground state) or serum/LIF media with mTOR inhibitor (mTORi, paused state). The 2i-induced ground state resembles naïve pre-implantation E4.5 epiblast cells (Ying *et al*, 2008; Ghimire *et al*, 2018). Global rewiring of signaling, metabolism and epigenome have been observed in the SL-2i transition. CpG methylation is dramatically decreased genome-wide, concomitant with a broad increase in H3K27me3 (Walter *et al*, 2016; Kumar & Elsässer, 2019). Further, a reduction of global H3K4me3 levels results in diminished promoter bivalency of developmentally regulated genes, where H3K27me3 and H3K4me3 are thought to set up a poised state (Sachs *et al*, 2013; Marks *et al*, 2012; Kumar & Elsässer, 2019; Atlasi & Stunnenberg, 2017). Enhancer activity is rewired during the SL-2i transition via Esrrb binding and H3K27ac activation (Atlasi *et al*, 2019). mTOR inhibition suppresses cell growth and division while retaining pluripotency, resembling the diapaused blastocysts *in vivo (Bulut-Karslioglu et al, 2016)*.

While most studies have focused on rewiring of regulatory circuits, activating or disengaging individual enhancers and transcripts, 2i and mTORi states also exhibit dramatically lower total cellular mRNA and attenuated transcription activity revealed by cell-number-normalized RNA-seq and EU (5-Ethynyl Uridine) incorporation (Bulut-Karslioglu *et al*, 2016). Given that transcript levels are balanced by rates of RNA synthesis and degradation (Schwalb *et al*, 2016; Herzog *et al*, 2017), to which extent transcription activity itself influences transcript abundance remains to be addressed. Quantitative RNA labeling techniques enable the absolute measurements of either RNA synthesis or degradation (Schwalb *et al*, 2016; Schwanhäusser *et al*, 2011; Rabani *et al*, 2011; Herzog *et al*, 2017; Muhar *et al*, 2018; Schofield *et al*, 2018). To uncover the transcription and chromatin interplay in SL-2i and SL-mTORi transitions, we measured both total and nascent RNA with TT-seq and estimated RNA turnover using spike-in references. We *de novo* annotated known and new transcription units (TUs) and examined how transcription of different TUs responds to state transitions. Our data reveals how the balance of RNA synthesis and dilution by cell growth/division shapes total RNA abundance in different pluripotent states. We also estimate RNA polymerase (Pol) II elongation velocity, characterize its manifestation in epigenomic features, and validate its impact on termination read-through distance.

## Results

### Transcription unit annotation in mESC pluripotent states

To capture newly synthesized RNA in different pluripotent states and transitions, we switched ES cells from SL medium to 2i medium for 2 or 7 days, mTORi medium for 1 or 2 days, pulsed 4sU (4-thiouridine) for 5 minutes and performed the TT-seq method as previously described (Schwalb *et al*, 2016) (Fig 1A). In order to verify our nascent RNA sequencing results, we compared gene coverage with GRO-seq (global run-on sequencing) (Flynn *et al*, 2016; Wang *et al*, 2015), PRO-seq (precision run-on sequencing) (Engreitz *et al*, 2016; Lloret-Llinares *et al*, 2018), NET-seq (native elongating transcript sequencing) (Mylonas & Tessarz, 2018; Tuck *et al*, 2018), 4sU-seq (Benabdallah *et al*, 2019; Brown *et al*, 2017) and Bru-seq (bromouridine sequencing) (Ardehali *et al*, 2017). Due to the short labeling time and fragmentation step included in the protocol, TT-seq nascent RNA showed a steadt coverage over the gene body, balanced signal from the first to the last exons as well as the highest intron/exon coverage ratio (Fig 1B, Fig EV1A-B). Thus, a stranded TT-seq signal provides an excellent demarcation of the nascent transcription unit and polymerase-independent measure of transcriptional activity.

**Figure 1.**
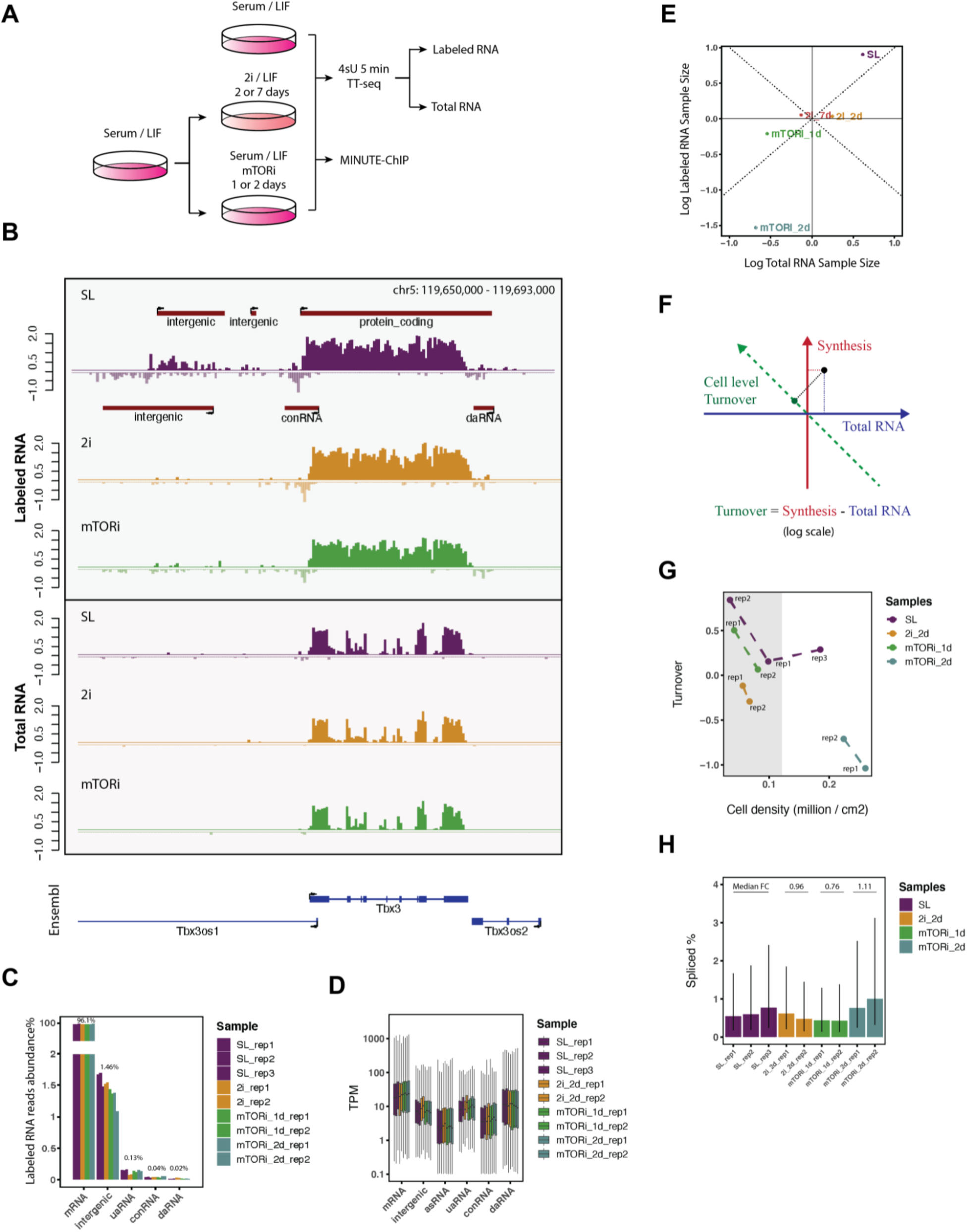
Mouse ES cells transcript annotation with TT-seq. A. Experimental Scheme for collecting TT-seq samples in Serum/LIF (SL), 2i/LIF (2i), Serum/LIF/mTORi (mTORi) conditions. B. TT-seq nascent RNA and total RNA profiles of gene Tbx3, with annotated transcription units (TU, red) and Ensembl gene references (blue) in mm10. C. Labeled RNA reads distribution by TU locations in percentage (y axis breaks from 2 to 40%). D. Labeled RNA TPM (transcript per million) by TU locations of each sample and replicate. E. Labeled and total RNA sample sizes distribution of the mESC pluripotent states. Sample sizes are represented by the log-transformed DESeq’s size factors after spike-in normalization. F. A schematic diagram of cell level RNA turnover approximation by the contrast of labeled and total size factors, which equals the projection on the green diagonal dashed line. G. Cell level turnover rates plotted against culture densities at time of harvesting. Replicates falling into the grey area of similar cell density were used to calculate spike-in normalized transcript abundance in Fig 2. H. Proportions of spliced labeled RNA reads (CIGAR ‘N’) on Ensembl protein-coding genes. The error bar shows the (0.25, 0.75) quantile of spliced ratio.

The pluripotent ES cell genome is pervasively transcribed (Efroni *et al*, 2008). We used nascent TT-seq signal to *de novo* define coding and non-coding TUs in the pluripotent genome separately for each condition and replicate. An R shiny application (TU filter) was developed offering previously described algorithms for TU discovery (Schwalb *et al*, 2016) within a simple and reproducible visual workflow (Materials and Methods). Overall, 96% of the uniquely mapped TT-seq nascent RNA reads could be assigned to 11743 GENCODE protein-coding genes, while approximately 1.5% were assigned to 20437 new intergenic RNAs called from TT-seq (Fig 1C). We automatically annotated TUs by their location and direction relative to transcription start site (TSS) or transcription termination site (TTS) of protein-coding TUs: upstream antisense (uaRNA), convergent (conRNA), *cis*-antisense (asRNA), downstream sense (dsRNA), downstream antisense (daRNA) TUs (Fig EV2A). An example of the TU annotation is shown for the Tbx3 gene neighborhood (Fig 1B). The sensitivity of our TU discovery pipeline is majorly determined by sequencing depth (Fig EV1C), thus accuracy of TU calling with TU filter relies on continuous read coverage at sufficient depth. Transcription of intergenic TUs was generally lower, suggesting that their cryptic/non-canonical promoters are generally weaker than divergent gene promoters (Fig 1C). Non-coding RNAs synthesized from the opposite strand of known genes were lowly transcribed (Fig 1D). Thus, lower expressed classes of intergenic RNAs generally exhibited higher inconsistency between replicates (Fig EV2C). uaRNAs, commonly arising from divergent transcription initiation at gene promoters, were most robustly called and relatively abundant amongst non-coding nascent transcripts (Fig 1C, Fig EV1C).

We also applied the same annotation process to published GRO-seq and PRO-seq to test if lowly expressed TUs were recapitulated across methods. TT-seq identified a lower number of non-annotated and method-specific short ncRNAs compared to GRO-seq and PRO-seq, reflecting the different labeling preferences of metabolic and run-on labeling approaches (Fig EV1D-F). Using ChIP-Seq profiles of RNA polymerase (Pol) subunits (Materials and Methods), TUs could be assigned to Pol I, II or III transcripts, albeit in many cases available occupancy profiles were insufficient to unequivocally link the TU to only one of the polymerases (Fig EV1G-H; Materials and Methods). Intergenic RNAs commonly called with all three methods showed the highest proportion of FANTOM5 enhancers (Andersson *et al*, 2014), ATAC-seq peaks (Atlasi *et al*, 2019) and STARR-seq enhancers (Peng *et al*, 2020), and showed low ChIP co-occupancy of all three RNA polymerase types (Fig EV1I). These results suggest that the majority of intergenic RNAs in mESCs exhibit low RNA polymerase occupancy and transcription frequency.

### Attenuated RNA synthesis in 2i and mTORi conditions

Next, we assessed transcriptional changes associated with 2i and mTORi transitions using TT-seq nascent RNA measurements. We used spike-ins to scale labeled and total RNA (Fig 1E; Materials and Methods) and confirmed that both transcription and total RNA abundance decreased in 2i and mTORi states (Bulut-Karslioglu *et al*, 2016). We approximated cell-level RNA turnover as a function of synthesis and total RNA measurements (Fig 1F) and observed a decrease in global RNA turnover in 2i and mTORi (Fig 1G). Cell-level RNA turnover is influenced by dilution via cell division, and the observed decrease in 2i and mTORi is in line with a slow-down of cell cycle progression in the new states (Bulut-Karslioglu *et al*, 2016). Accordingly, we also observed a downward trend in turnover rate as cell confluency increased (Fig 1G). To exclude the possibility that the different states would have intrinsically different labeling efficiencies, we confirmed similar fractions of spliced nascent transcripts across the three states (Fig 1H) (Wachutka *et al*, 2019).

### Genome context modulation underlies global transcription variation

Next, we counted the labeled RNA reads on mRNA, intergenic RNA, uaRNA and asRNA intervals, and compared to 2i, mTORi to SL conditions using spike-in normalization (Fig 2A). Transitioning from SL to mTORi was associated with a homogeneous down-regulation of both mRNA and ncRNA synthesis, while individual mRNAs and intergenic RNAs showed a highly variable response with both up- and down-regulation in 2i. mRNA synthesis remained largely correlated between SL in mTORi condition, irrespective of initial expression level, while intergenic transcripts were more variable (Fig 2B, right). In contrast, 2i induced a greater extent of rewiring for both mRNAs and ncRNAs (Fig 2B, left). Intergenic TUs in general followed the global decrease in transcription in 2i and mTORi conditions (Fig 2C). A notable exception were bidirectional TUs lacking clear enhancer status, which were up-regulated in 2i condition relative to the global downward trend (Fig 2A, C). From a chromatin perspective, H3K27me3-repressed/bivalent ChromHMM states exhibited larger coefficients of predicting log2 fold-change for bidirectional than unidirectional intergenic TUs (Fig EV2D). This may indicate low connectivity of spurious intergenic transcription units with nearby genes, and imply a basal suppression through H3K27me3 (Fig EV2E) (Haberle & Stark, 2018).

**Figure 2.**
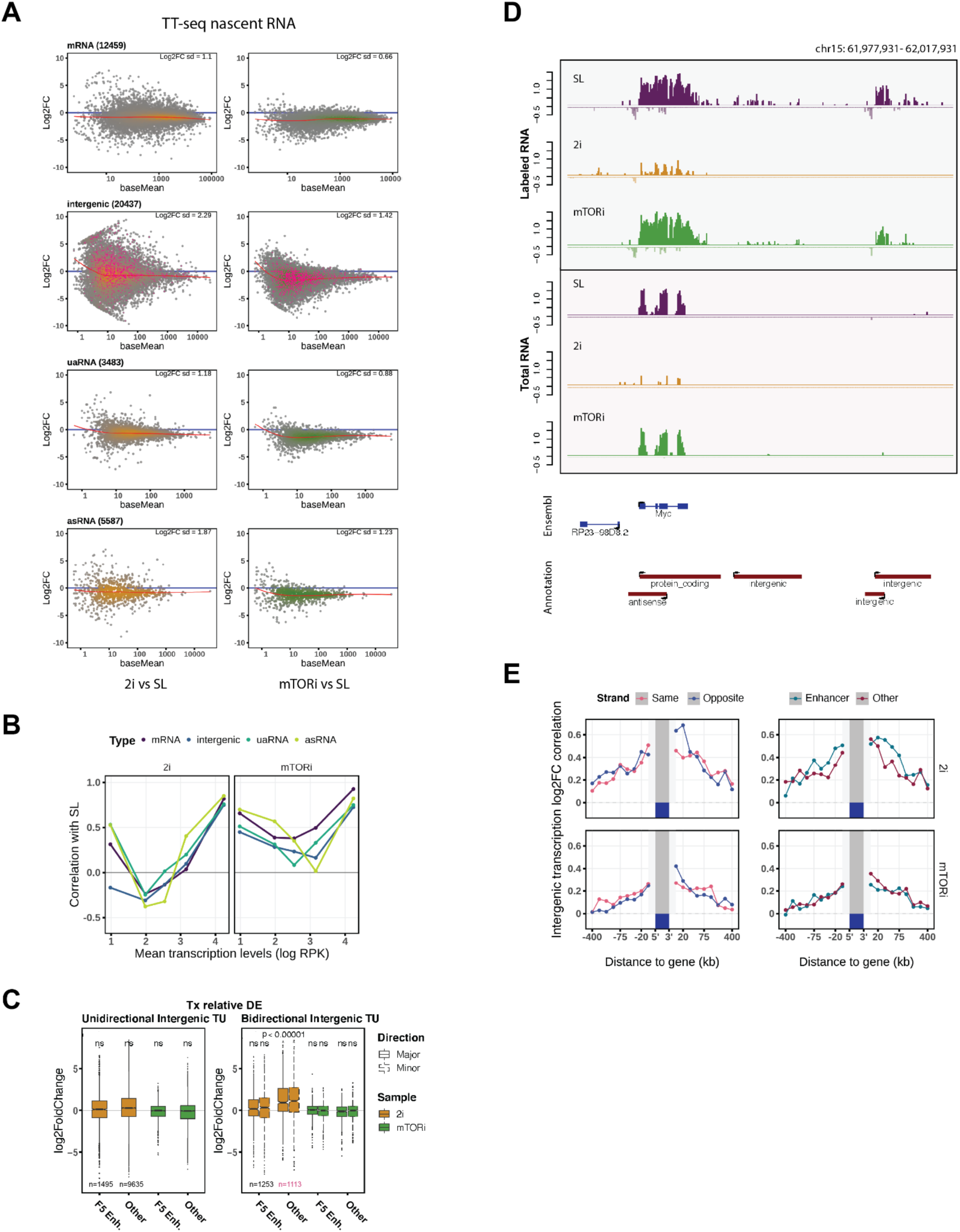
Pluripotent states transcription rewiring. A. Spike-in normalized MA-plots in the contrasts of 2i (2 days, yellow) and mTORi (1 day, green) to SL state. Local regression lines are in red. Bidirectional non-enhancer intergenic TUs in C are highlighted in pink. B. Pearson’s correlation of log labeled RNA RPK by mean transcription level bins in the same contrasts. C. DESeq2 internal normalized differential expression tests of intergenic RNA labeled RNA read counts. Major and minor TU directions were assigned by labeled RNA RPKs. FANTOM5 (Andersson *et al*, 2014) mouse enhancers were overlapped with the intergenic TUs. Statistical test was performed with two-tailed Student’s t-test of log2 fold changes. D. Spike-ins normalized labeled and total RNA profiles of Myc gene neighborhood E. Log2FC correlation of neighboring intergenic TUs and genes by genomic distance bins, separately for the relative strandedness to genes and the combined enhancer annotations (FANTOM5(Andersson *et al*, 2014), ChromHMM(Pintacuda *et al*, 2017) and STARR-seq(Peng *et al*, 2020)).

To assess the contribution of RNA synthesis to changes in total RNA abundance, we compared the pairwise correlation of the three states’ labeled RNA and total RNA length-normalized read counts (log-RPK) (Fig EV2G). mRNA, intergenic RNA and asRNA labeled RNA levels correlated well with their total RNA levels (Fig EV2F), which suggests their total RNA abundance is dominantly regulated via RNA synthesis, and to a lesser extent by degradation, across the three states. This is in line with recent studies demonstrating that rewiring of promoter-enhancer interactions, thus changes in transcriptional activity, explains changes in RNA-seq abundance between SL and 2i states for many differentially regulated genes (Atlasi *et al*, 2019; Joshi *et al*, 2015; Novo *et al*, 2018).

We wondered whether transcriptional changes of intergenic TUs followed a discernable mechanism. Myc gene expression is known to be strongly attenuated under 2i treatment (Marks *et al*, 2012) (Galonska *et al*, 2015) but is maintained in mTORi (Bulut-Karslioglu *et al*, 2016), and interestingly, we found the same trends to govern neighboring intergenic TUs (Fig 2D). Globally, intergenic TU transcription changes correlated well with transcription changes of neighboring genes (n=11684) in a direction- and proximity-dependent manner (Fig 2E). The highest correlation was observed for transcription changes within the 20 kb region downstream antisense, thus convergent transcripts of protein-coding genes (Fig 2E, top). Transcriptional changes at protein-coding genes correlated better with transcriptional changes at intergenic TUs overlapping an enhancer state than at other intergenic TUs within ±100 kb region of the gene (Fig 2E, top-right, Fig EV2H-I). We further validated the observed position regulated gene expression trend in a single-cell RNA-seq dataset (Kolodziejczyk *et al*, 2015) (Fig EV2J). In summary, our analysis supports a previously reported enhancer-gene position dependent co-regulation of the transcriptome during the SL-2i transition (Atlasi *et al*, 2019), while mTORi largely preserves relative mRNA levels and therefore exhibits less correlated changes between coding and intergenic transcripts (Fig 2E, bottom).

### Estimating elongation velocity with TT-seq and Pol II coverages

TT-seq nascent RNA coverage allows estimation of *productive transcription initiation frequencies* (Gressel *et al*), which is defined as the number of Pol II molecules that initiated at the promoter and were successfully released into productive elongation per unit of time. Meanwhile, Pol II ChIP-seq captures the Pol II occupancy on the DNA template in steady state, which depends on the number of polymerases and their elongation velocity (Ehrensberger *et al*, 2013). Thus, by combining Pol II occupancy measurements with TT-seq derived initiation frequencies, Pol II elongation velocities can be derived (Caizzi *et al*, 2021). Since Pol II S5p (CTD Serine-5 phosphorylation) appears at TSS after the pre-initiation complex dissociation (Søgaard & Svejstrup, 2007; Glover-Cutter *et al*, 2009), its signal corresponds most closely to the fraction of productive Pol II amongst the total chromatin-engaged Pol II molecules (Steurer *et al*, 2018). Therefore, we used the ratio of TT-seq density over Pol II S5p density from quantitative, native MINUTE-ChIP collected under the same three conditions, SL, 2i and mTORi as a proxy for relative transcription elongation velocity (Materials and Methods). TT-seq nascent RNA / Pol II S5p ratio exhibited a deep minimum of estimated transcription velocity after the TSS, indicating promoter proximal pausing (Steurer *et al*, 2018) (Muse *et al*, 2007) (Bartman *et al*, 2019); a flat gene body ratio demonstrated the steady elongation velocity, and our transcription velocity estimation also captured the “getting up to speed” model towards elongation termination (Jonkers *et al*, 2014; Jonkers & Lis, 2015) (Fig 3A-C).

**Figure 3.**
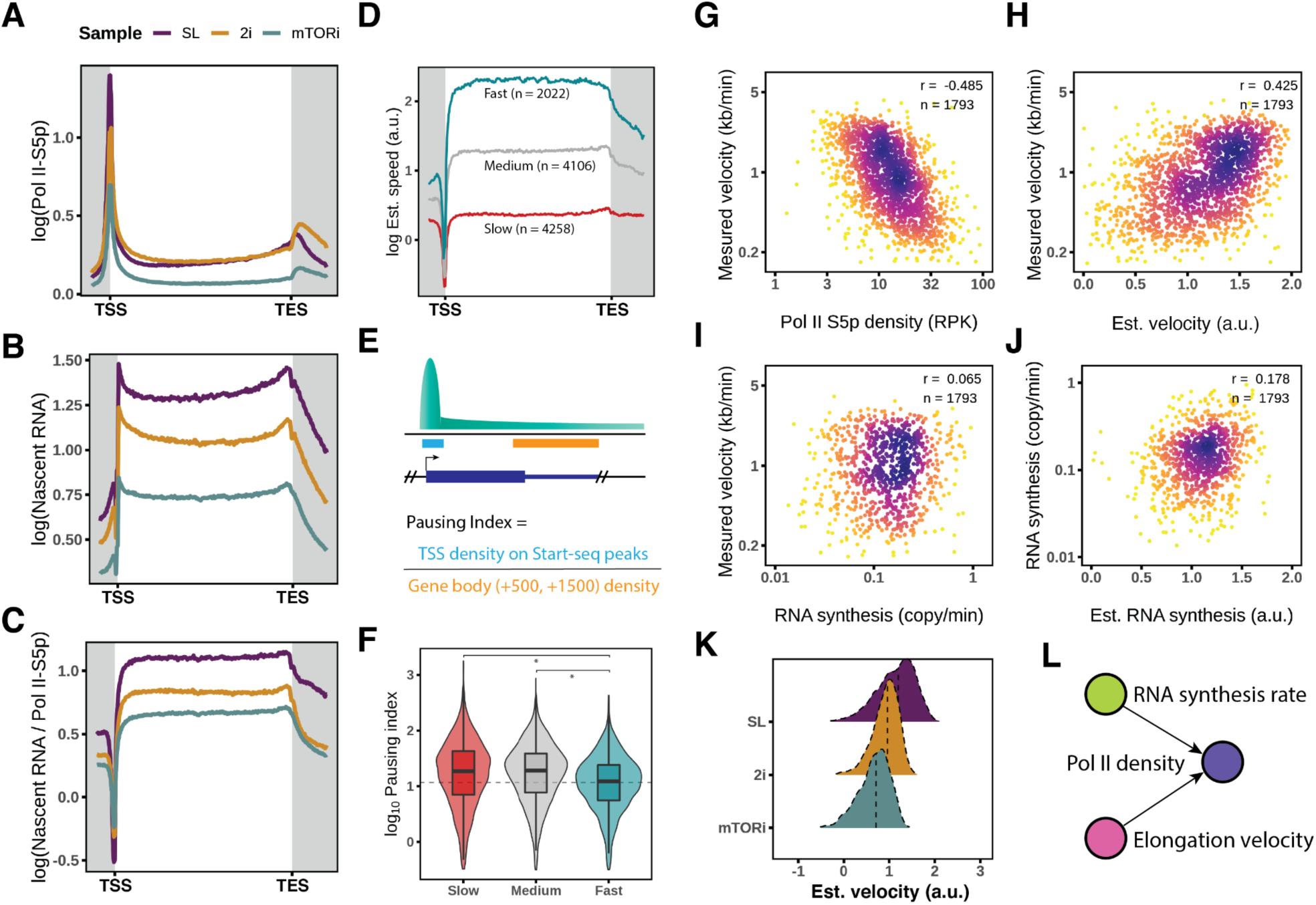
Elongation velocity estimation in the pluripotent states. A-C. Quantitative MINUTE-ChIP mean coverage profiles of Pol II S5p, spike-ins normalized TT-seq nascent RNA and their ratio as the estimated elongation velocity. D. K-means grouped average profiles of estimated velocity in the SL condition, the same condition for F-J. E. Illustration of pausing index calculation (Materials and Methods). F. Pausing indexes of Pol II S5p by different elongation velocity groups in the SL condition (* Student’s t-test p < 2.2e-16). G. Correlation of measured elongation velocity (Jonkers *et al*, 2014)(Materials and Methods) and Pol II S5p gene body occupancy. H. Correlation of GRO-seq measured velocity and estimated velocity by TT-seq nascent RNA and Pol II S5p ChIP ratio. I. Correlation of measured velocity (Jonkers *et al*, 2014) and TT-seq measured RNA synthesis rates. J. Correlation of TT-seq measured RNA synthesis rates and estimated RNA synthesis rate by Pol II S5p density and GRO-seq measured velocity (Jonkers *et al*, 2014). K. Ridge distribution of estimated elongation velocity of the pluripotent states. L. A scheme describing the relationship of RNA synthesis rate, elongation velocity and Pol II gene occupancy.

Next, we closely examined the estimated velocity in SL, and used k-means to cluster the estimated elongation velocity profiles of 10386 protein-coding genes into three groups with slow, medium and fast elongation velocity (Fig 3D, Fig EV3C). Acceleration of elongation velocity towards the TES (transcript end site) was only observed for the slow and medium elongation velocity gene groups (Fig 3D). Beyond the TES, the fast elongation group showed a steeper decline of transcription velocity (Fig 3D, Fig EV3C). Since the genes in the slow and medium elongation velocity groups showed a deeper drop near the TSS, we wondered if this feature was indicative of increased Pol II promoter-proximal pausing. We thus verified this observation by calculating a pausing index (Fig 3F, Materials and Methods) and established that promoter-proximal pausing was least prevalent in the fast elongating gene group (Fig 3E).

To validate our velocity estimates, we recalculated elongation velocities from Cdk9 inhibition (Cdk9i) time course GRO-seq experiment (Jonkers *et al*, 2014) (Materials and Methods). The recalculated velocities agreed with the published velocities but included a larger number of genes (Fig EV4A). Comparing our estimated elongation velocities to the Cdk9i-derived direct elongation velocities revealed a good global correlation (Fig 3H, Pearson’s r=0.425). Thus, the labeled RNA / Pol II S5p ratio provides a general way of estimating elongation velocity transcriptome-wide. In addition, the estimated velocity reversely correlated with Pol II S5p density (Fig 3G, r = −0.485) but not RNA synthesis rates (Fig 3I, r=0.065). Therefore, with the assumption that Pol II coverage results from the transcription initiation frequency (RNA synthesis rate) and the elongation velocity (Fig 3L), we reversely estimated RNA synthesis rates from the velocity measured by CDK9 inhibition and Pol II S5p density, and these also agreed with a modest correlation with the actual RNA synthesis rates from our TT-seq data (Fig 3J, r=0.178). These results manifest the close connectivity of Pol II density with its two determinants, RNA synthesis rate and elongation velocity (Fig 3G, Fig EV3A), while RNA synthesis rate and elongation velocity are independently controlled. Therefore, combining Pol II occupancy and RNA synthesis rate is also able to inform on the local elongation velocity.

### Interpretation of elongation velocity in mouse ES cells

We compared elongation velocity in 2i and mTORi cells, and found 2.2 and 3.7 fold decrease in median elongation velocity relative to SL cells (Fig 3K). 2i cells showed a preferential reduction of fast-elongated genes, while mTORi cells slowed down homogeneously (Fig EV4B). To understand the regulation of elongation velocity, we first sought to correlate elongation velocity in SL condition with other epigenomic features. Many active histone modifications have been found to be associated with elongation velocity (Jonkers *et al*, 2014; Veloso *et al*, 2014), for instance H3K36me3 and H3K79me2. Using our estimation, we confirmed that these two active marks positively correlated with elongation velocity and identified a positive correlation with several chromatin remodelers (Chd1, Chd2 and Chd9) (Fig 4A). Notably, a recent Cryo-EM structure implicates Chd1 in clearing chromatin in front of the traversing RNA polymerase (Farnung *et al*, 2021). Moreover, we observed anti-correlation with Polycomb repressive marks (H3K27me3, H2Aub and Ezh2) histone variant H2A.Z, also shown to negatively associate with pause release (Mylonas *et al*, 2021) (Fig 4A).

**Figure 4.**
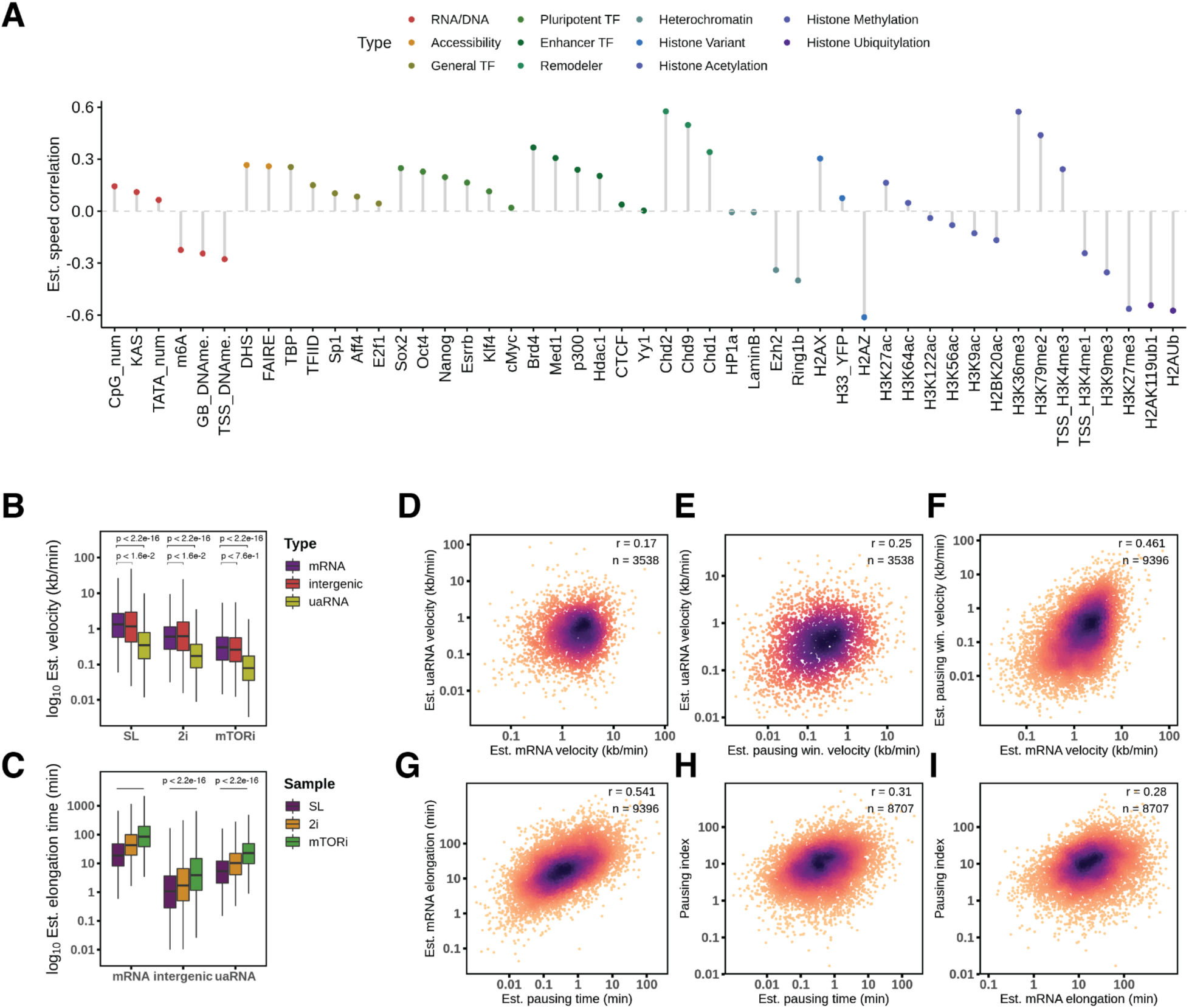
Transcription elongation velocity interpretation. A. Correlation of estimated gene elongation velocity (n = 10611) with sequence and chromatin features (Materials and Methods). B. Estimated velocity scaled by GRO-seq measured velocity plotted across the culture conditions and TU types. p-values were tested with Student’s t-test in log scale. C. Estimated velocity scaled by GRO-seq measured velocity plotted across the culture conditions and TU types and estimated elongation time from the respective TU lengths. P-values were tested with Student’s t-test in log scale. D-F. Estimated velocity correlation between mRNA, paired uaRNA and mRNA TSS pausing window by Start-seq. Estimated elongation dynamic parameters were in SL condition, the same as below. G-H. Correlation of estimated pausing duration on Start-seq TSS intervals with gene body elongation time and pausing index. I. Correlation between estimated mRNA gene body elongation time and pausing index.

We observe Pol II to travel faster through mRNA and intergenic TUs than uaRNA TUs (Fig 4B). Deriving TU elongation time estimates as elongation velocity divided by TU interval length, we found that Pol II spends the longest time traversing through mRNAs, followed by uaRNAs and intergenic TUs (Fig 4C). When looking at pairs of uaRNA and mRNA arising from a shared promoter region, we found a moderate correlation in estimated elongation velocity and time (Fig 4D-F).

Next we questioned if Pol II pause-release dynamics were connected to elongation velocity or RNA synthesis rate. Transcription initiation frequency is known to weakly anti-correlate with promoter-proximal pausing (Sanchez *et al*, 2018; Gressel *et al*, 2019; Gressel 2017). Accordingly, we observed this trend consistently in all the three conditions (Fig EV3B). The average pause duration, which corresponds to the time Pol II needs to travel through the promoter-proximal pausing interval, has been found to be 42 seconds in human fibroblasts (Steurer *et al*, 2018). We approximated and scaled the pause duration with the external scale of GRO-seq measured elongation velocity, and agreed that the median pause durations were 21, 37 and 61 seconds for SL, 2i and mTORi cells, respectively (Fig EV4C). Then we examined pause durations in SL and found a good correlation with gene elongation time (Fig 4G). Further, we also observed that the pausing index moderately correlated with pausing time and mRNA elongation time (Fig 4H-I). These results indicate that RNA polymerase II pause-release dynamics connects with pausing and elongation duration, while gene body elongation time associates with TSS pausing duration to a larger extent (Fig 4G).

### Transcription termination read-through is associated with elongation velocity

In K562 cells, TT-seq mapped the ultimate termination sites of (C/G)(2–6)A motifs enrichment around 3kb down-stream from the last pA (poly-adenylation) sites (Schwalb *et al*, 2016). To define genome-wide ultimate termination sites from our TT-Seq data we used a method to determine the maximal density contrast (Fig 5A, Materials and Methods). Next we applied the same algorithm to other transcription readout methods (PRO-seq and Pol II S5p ChIP-Seq), and found that the termination windows agreed with TT-seq results in SL and 2i conditions, which suggests an inherent termination mechanism (Fig EV5A).

**Figure 5.**
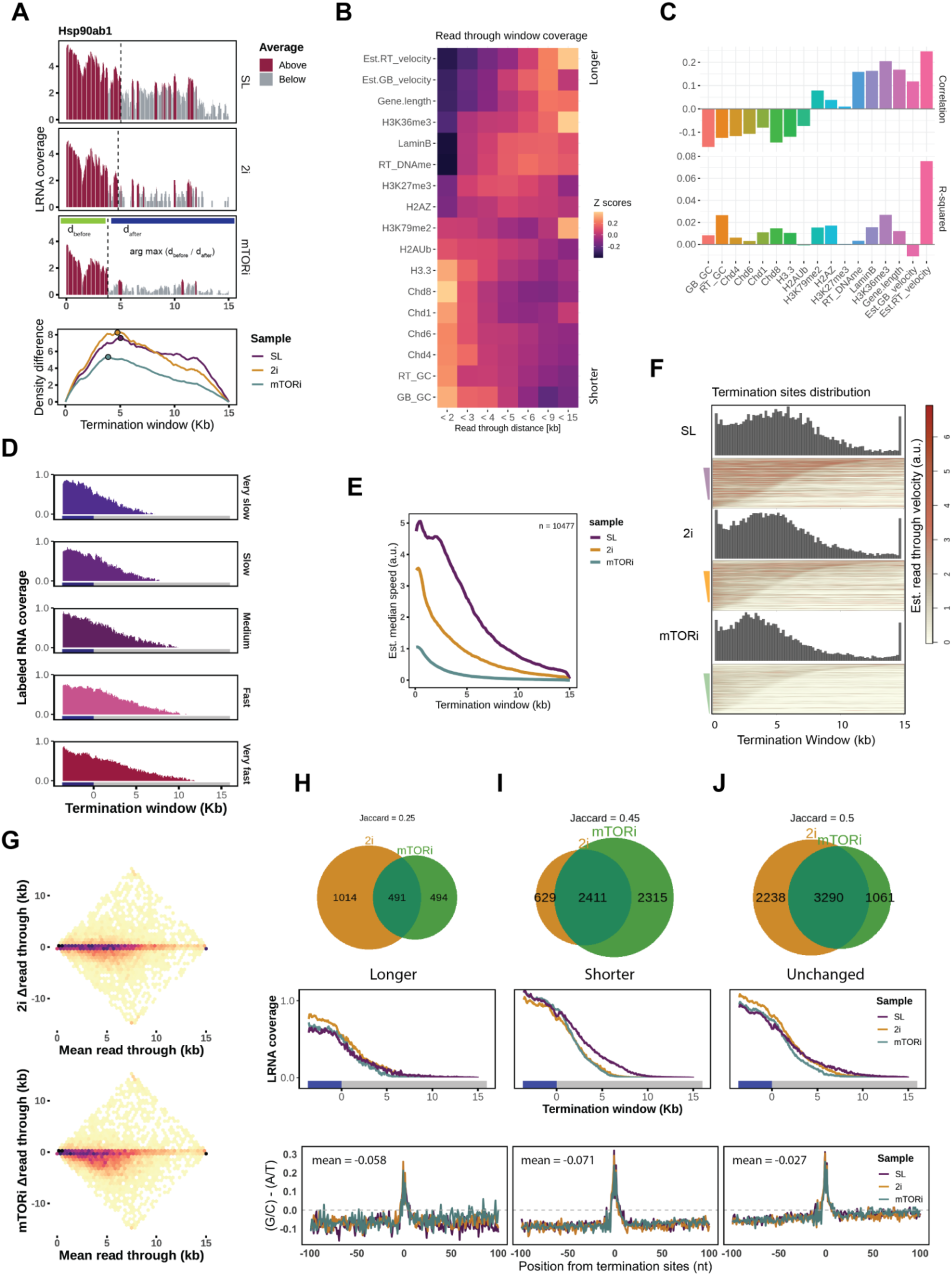
Transcription termination read-through with elongation velocity. A. Illustration of termination site detection algorithm on gene Hsp90ab1, by maximizing the read density contrast as shown in the sliding curves (bottom) (Materials and Methods). The ultimate termination sites were indicated by dash lines and solid points for the three cell states. B. Gene features and chromatin features termination window occupancy by binned read-through distances. Values were log normal transformed and standardized to z-scores. C. Correlation of read-through distances with individual chromatin features (upper); multivariate linear regression decomposed R-squared values for the read-through distance explanation (lower) (Materials and Methods). D. Labeled RNA median coverage in the termination window ordered by measured elongation velocity (Jonkers *et al*, 2014). E. Estimated elongation velocity median coverage in the termination window by the mESC states. F. Termination sites distribution (above) and estimated elongation velocity coverage heatmaps (below) by the order of read through distance in each condition. G. Scatter plots of read-though distance changes in 2i and mTORi states with mean read-through distance. H-J. Classes of read through distance changes in 2i and mTORi state. Top, Venn diagrams with Jaccard overlapping index; middle, labeled RNA reads coverage from the last exon (blue box) to 15 kb downstream termination window (grey box); bottom, GC versus AT nucleotide mean frequency contrast in the 100 bp flanking region around termination sites.

Analysis of Pol II mutants revealed that mutants with slower elongation velocity also exhibited a shorter termination window (Hazelbaker *et al*, 2013; Sheridan *et al*, 2019). To which extent the natural elongation velocity explains termination read-through has not been determined. Therefore, we analysed TT-seq nascent RNA coverage in a 15 kb termination window across groups of genes with increasing elongation velocity and found that indeed fast elongation genes had longer read-through distances (Fig 5D).

We wondered if elongation velocity was the dominant determinant of read-through distance. To this end, we calculated the average velocity in the read-through window of each gene and evaluated the extent of which elongation velocity versus a large panel of known epigenomic features can explain the read-through distance. We first assumed linear responses and ranked with Pearson’s correlation. The long read-through distances (>9 kb) associated with the faster estimated read-through/gene body velocity, higher H3K36me3 and gene lengths. In contrast short read-through (<3 kb) windows associated with H3.3, Chd8, Chd1 enrichment and GC content in the termination window (Fig 5B). A total of 25% read-through distance variance could be explained by a multivariate linear model, in which the estimated read-through velocity explained the largest variance of read-through distance (Fig 5C). We further confirmed read-through velocity’s importance with a gradient boosting machine (gbm) nonlinear model, which explained 56% variance with acceptable prediction accuracy of the read-through distances (Fig EV5C-E). At the ultimate termination sites (−1 kb, 1kb) windows, we also found co-localization of chromatin accessibility and daRNA initiation, which suggest antisense transcription collision may participate in the termination process for a small number of cases (Fig EV5B).

### Attenuated elongation velocity associates with shorter read-through distance

By our estimated termination sites, slower elongation in 2i/mTORi coincided with shorter median read-through distances: SL 5.1 kb, 2i 4.6 kb and mTORi 3.9 kb (Fig 5F). Termination sites in 2i and mTORi cells were mainly located within 5 kb downstream of the transcript end site (TES) (Fig 5F). In line with this, the estimated elongation velocity rapidly declined in 2i/mTORi cells within 5 kb after the TES, while in SL condition median estimated elongation velocity decreased more gradually (Fig 5E). We noted that the steepest decline in elongation velocity immediately preceded the ultimate termination site (Fig 5F). Thus, our data supports a model in which Pol II encounters termination roadblocks imposed by DNA sequence, and the remaining velocity at the encounter stochastically determines if the polymerase comes to a complete stop.

To understand if altered elongation velocity in 2i and mTORi conditions also affected termination read-through distance, we grouped genes according to their read-through distance shortening, extending or remaining unchanged with ±500 bp (Fig 5G-J). Genes with an extended read-through distance in 2i or mTORi were not associated with a clear extension of the average TT-seq nascent signal and represented sporadic cases where relatively low labeled RNA read coverage may have prohibited accurate termination site determination (Fig 5H). Genes with a shortened read-through distance in 2i or mTORi also exhibited a marked drop in RNA synthesis, where labeled RNA coverage reached a baseline several kb earlier (Fig 5I).

We extracted the GC nucleotide frequency around the termination sites for the three different groups. Irrespective if the termination shifted to an earlier or later site, both the old and new termination sites exhibited a sharp local maximum in G/C-richness (Fig 5I-J). This recapitulated the G/C-rich motif mapped at K562 cells termination sites (Schwalb *et al*, 2016). Therefore, switching pluripotent conditions, the main termination of genes either remained centered on the same G/C-rich sequence motif, or switched to an earlier or later G/C-rich sequence motif. In the model described above, G/C-rich motifs thus provide roadblocks to RNA polymerase II that lead to a final termination if the velocity is already sufficiently reduced.

## Discussion

With the assistance of quantitative techniques, our data uncovers global transcriptional features distinguishing related pluripotent states of mouse embryonic stem cells. Our data supports a model in which inhibition of MAPK/GSK3b or mTOR signaling pathways by 2i or mTORi, respectively, directly affect global transcriptional output and RNA polymerase elongation dynamics without proportional changes in chromatin features known to associate with protein-coding gene transcription initiation or elongation.

Of note, inhibition of mTOR signaling homogeneously reduced RNA synthesis rates to a large extent and only marginally redistributed transcriptional activity on both coding and non-coding TUs (Fig 1B-C; Fig EV2C). In contrast, the SL-to-2i transition leads to a larger rewiring of mRNA gene expression. Intriguingly, transcriptional changes occur mostly without changes in enhancer-promoter interactions (Atlasi *et al*, 2019; McLaughlin *et al*, 2019), suggesting that enhancer-promoter pairing provides hardwired connectivity that remains responsive to cellular signaling (Schoenfelder & Fraser, 2019). 2i-specific enhancer activation occurs via the loading of Esrrb upon the pre-existing chromatin interactions (Atlasi *et al*, 2019), resulting in H3K27ac enrichment (Schoenfelder & Fraser, 2019; Barakat *et al*, 2018), which itself appears characteristic albeit not necessary for enhancer function (Zhang *et al*, 2020; Sanchez *et al*, 2018). In our data, intergenic ncRNAs showed a strand and distance dependent co-regulation with neighboring coding genes, which was associated with enhancer potentials in 2i cells but not in mTORi cells (Fig 2E). The global rewiring of cell signaling and metabolism in the ground state, therefore, reveals the large extent of co-regulation of ncRNAs with their adjacent genes (Engreitz *et al*, 2016) (Fig 2F, Fig EV2H). Spurious transcription, such as bidirectional intergenic TUs without enhancer annotation or active chromatin states do not follow the global transcriptional repression in 2i (Fig 2C), which may be an effect of the global CpG hypermethylation observed in 2i (Jin *et al*, 2017; Haberle & Stark, 2018; Walter *et al*, 2016).

Previous estimates of transcription elongation kinetics rely on chemical inhibition and time-series experiments, either by transcription release after the inhibitor clearance (Danko *et al*, 2013; Fuchs *et al*, 2014; Veloso *et al*, 2014), Pol II initiation/pause-release repression (Jonkers *et al*, 2014) or RNA reporter elongation efficacy (Fukaya *et al*, 2017). Such velocity estimations still leave details of global and local elongation dynamics unaddressed (Jonkers & Lis, 2015). Furthermore, it has been shown that with a multi-omics approach, which combines TT-seq with mNET-seq derived Pol II occupancies, transcription elongation velocities can be estimated (Caizzi *et al*, 2021). Accordingly, we used here the ratio of TT-seq nascent RNA and Pol II S5p coverage as an approximation of elongation velocity and show that it provides a universal and transcriptome-wide measurement of several key parameters: First, we were able to estimate the unperturbed elongation velocity coverage continuously from initiation to termination (Fig EV3C). Second, we were able to compare native elongation velocity across different cell states (Fig 3C). Third, we deduced the relationship of elongation velocity, RNA synthesis and Pol II occupancy, which supports the observation that elongation velocity has a minimal impact on RNA synthesis (Jonkers *et al*, 2014). Fourth, we mapped the local transcription dynamics downstream of the TSS and in the termination window, and recapitulated the relationships of pausing/elongation duration and elongation velocity/read-through distance. Although low gene expression impedes local velocity estimation from TT-seq nascent RNA and Pol II coverages, a TU-level velocity estimation, averaging TT-seq signal on the entire TU, provides a reliable comparison across conditions. Therefore we were able to deduce that RNA synthesis decrease in the pluripotent state transitions is independent from elongation slowdown and neither controlled by pause release dynamics (Fig 3L, Fig EV3B). This result agrees with a recent study that transcription initiation dominates gene regulation in murine erythropoiesis differentiation over pause-release dynamics (Larke *et al*, 2021).

Finally, elongation velocity explains the termination read-though length more than any other chromatin feature, although the ultimate elongation slow-down may manifest Pol II disassembly by the RNA cleavage-mediated “torpedo” model (West *et al*, 2004; Brannan *et al*, 2012; Nojima *et al*, 2015; Baejen *et al*, 2017) (Fig EV5B). Given the read-through distances were well preserved across sequencing methods (Fig EV5A), we can conclude that slower elongation in 2i and mTORi conditions brought only a fraction of genes to earlier termination with the majority of read-through distances inherently unchanged (Fig 5J, Fig EV5A). A higher frequency of read-through window shortening was observed in in mTORi cells. but the exact changes of termination window length are gene-specific and may depend on GC-rich motif occurrence in the termination window (Fig EV6A-B, Fig 5I). Nevertheless, the large overlap of genes with shorter read-through in 2i and mTORi cells reveals the subgroup of genes that are responsive to elongation velocity slowdown (Fig 5I-J). Therefore, at the cell population level, multiple termination sites may exist simultaneously, among which our method detects the most frequent site in use (Fig 5B). Elongation velocity reduces relatively quickly upon passing the TES (Fig 4D) but continues to traverse at a slower level until the final termination site. Thus, read-through velocity has a more direct influence on the termination site usage as compared to other chromatin features (Fig 5F).

In sum, our findings reveal transcription dynamic changes in the pluripotent states, and suggest the decisive role of RNA synthesis in regulation of coding and non-coding RNAs cellular abundance. Our analysis of elongation velocity with the multi-omics strategy provides new insights into promoter-proximal pausing and termination read-through in mouse ES cells.

## Materials and Methods

### Cell culture

Mouse embryonic stem cell RW4 (male, 129X1/SvJ) were cultured 0.1% gelatin-coated dishes with serum medium: Knock-out DMEM medium with 15% FBS (Sigma, F7524), 0.1mM ESGRO LIF (Sigma, ESG1107), 2 mM GlutaMAX (ThermoFisher, 10565018), 0.1 mM Non-Essential Amino Acid (Sigma, M7145), 0.1 mM β-mercaptoethanol (Sigma, M3148); 2i medium: ESGRO Complete Basal Medium (Millipore, SF002), 3 μM GSK3β inhibitor CHIR99021 (Sigma, SML1046), 1 μM Mek 1/2 inhibitor PD0325901 (Sigma, PZ0162), 0.1 mM LIF. Inhibition of mTOR was in serum-LIF (SL) medium supplemented with 200nM INK128 (CAYM11811-1).

### TT-seq extraction protocol

TT-seq labeling steps were performed as described before (Gressel *et al*, 2019; Schwalb *et al*, 2016) with minor modifications. In short, cells in the different pluripotent media were cultured for 1-2 days in four 15 cm dishes. With one dish for cell number counting, and the rest were supplemented with 500 µM of 4-thiouridine (4sU) (Sigma-Aldrich, T4509) for 5 min at 37°C and 5% CO2, then immediately quenched by adding TRIzol (ThermoFisher, 15596018) for RNA extraction. Total RNA was fragmented to 1kb with Bioruptor (Diagenode), then coupled with HPDP-Biotin (ThermoFisher, 21341) dissolved in dimethylformamide (VWR, 1.02937.0500). A small aliquot was saved as fragmented total RNA (FRNA) and the rest was purified with µMACS streptavidin beads (Miltenyi Biotec, 130-074-101). Purified labeled RNA (LRNA) and FRNA were subjected to DNase digestion (Qiagen, 79254). Total fragmented RNA and labeled RNA libraries were prepared with Ovation Universal RNA-Seq kit (NuGEN, 79254). The pooled library was size-selected by Ampure XP beads (Beckman Coulter, A63881) before sequencing with NextSeq® 500/550 High Output Kit v2 (Illumina, FC-404-2005, 75 cycles).

### Reads alignment and TU annotation

Paired-end short reads were aligned to mm9 and mm10 genome references (GENCODE) by STAR 2.7.3a with setting:

--outFilterMismatchNoverReadLmax 0.02 --outFilterMultimapScoreRange 0 --alignEndsType EndToEnd.

Mapped reads were subjected to transcription unit annotation as described before (Schwalb *et al*, 2016), processed in parallel by “TU filter” (R shiny app). Briefly, paired-end reads mid-points were binned into 200 bp genome coverage matrices by strand (merging the counts if sample has multiple replicates), then subjected to the binary hidden Markov state calling with R package “GenoSTAN” with “PoissonLog” method. Next the active state intervals were as the raw TUs and joined by exons per gene. Non-coding TU locations were named by their relative position to the nearby coding TUs (Fig EV2A).

TU differential expression analysis was performed by DESeq2 (1.24.0) (Love *et al*, 2014) with read counts on the annotated TU intervals by featureCounts (Rsubread 1.34.7) (Liao *et al*, 2019).

### Spike-in RNA design

ERCC synthetic spike-in RNAs were used as external reference for total RNA and labeled RNA sample size normalization as described in (Gressel *et al*, 2019; Schwalb *et al*, 2016) with minor modifications. Briefly, 6 pairs of spike-in RNAs with 4sU labeled/unlabeled mixture were prepared as below:

| ERCC spike-in RNA | Concentration (ng/μL) | Labeled rate (%) |
| --- | --- | --- |
| Sp2 (ERCC-00043) | 1 | 100 |
| Sp4 (ERCC-00136) | 0.1 | 100 |
| Sp5 (ERCC-00145) | 1 | 10 |
| Sp8 (ERCC-00092) | 0.1 | 10 |
| Sp9 (ERCC-00002) | 1 | 0 |
| Sp12 (ERCC-00170) | 0.1 | 0 |

For every million cells, 0.4 ng spike-in mix was added into the TRIzol (ThermoFisher, 15596018) cell lysis to eliminate technical errors retained during the steps of biotinylating, RNA purification and library preparation.

### Sample size estimation

We quantified GENCODE transcripts, de novo annotated TUs and spike-in transcripts using the alignment-free mapper (kallisto 0.46.2). For normalization, we calculated the size factors of all spike-ins in the total transcriptome and spike-ins 2-8 in the labeled transcriptome according to DESeq’s method (Love *et al*, 2014). The normalised transcriptomes were subjected to size factor calculation again and presented as the relative sample sizes of both total and labeled RNA abundance (Fig 1E).

### RNA synthesis rate estimation

The estimated labeled/total RNA read counts of GENCODE transcripts and spike-ins from kallisto were first normalized by respective spike-ins sizes from each sample. The highest expressed transcript of each gene was kept. A linear model was trained only with the normalized labeled spike-ins (Sp2, Sp4, Sp5 and Sp8) log2 labeled (X_L_) and total (X_F_) read counts in response to the respective label rates r: log_2_(r) ∼ X_F_ + X_L_. The evaluation was performed with 5-fold cross validation with 10 times subsampling, and the predictions on the hold-on set were examined. Then a final model with spike-ins counts of all samples were trained, which resulted in an adjusted R-squared 0.9917. And labeled rate prediction was made with transcript’s labeled/total read counts. In the same way, a second model was trained to predict all spike-ins weight per cell w: log2(w) ∼ X_F_ + X_L_, which resulted in an adjusted R-squared 0.991. Then the copy number per cell was transformed from weight with transcript effective length from kallisto. Finally, RNA synthesis rate (cell^-1^ minute^-1^, or copy/min per cell) was calculated by multiplying labeled rate and transcript copy number.

### Read coverage

Short reads genome coverage was processed with R/Bioconductor packages “rtracklayer” and “rsamtools” by piling up only the uniquely mapped and paired-end reads with insertion size less than 2 kb. Sample-wise coverage of each genomic interval was normalized by the respective size factors. For TT-seq sample size factors were generated from spike-in RNA read counts, for MINUTE-ChIP samples scaling factors were from the respective input sizes. In the heatmaps and coverage profile plots, each coverage vector of different lengths was resized by “spline” function to the same number of positions.

### Pol I, Pol II and Pol III TU classification

RNA polymerases ChIP-seq datasets ((Jiang *et al*, 2020), GSE145791) were aligned to mm10 genome by bowtie2 with “--local” setting. Each TU’s Pol I, Pol II and Pol III density were divided by the sum to obtain relative occupancies, and a threshold of 90% enrichment was used for the classification indicated in the ternary plot (Fig EV1H). For validation, ChIP-seq data of Pol III subunits PRC1, PRC4 and the cofactors BRF1, TFIII ((Carrière *et al*, 2012), E-MTAB-767) were aligned and subjected to peak calling with MACS2 default setting at “-q 0.01” cutoff, and overlapped on the ternary plot of the combined intergenic TUs (GRO-seq, PRO-seq and TT-seq).

### Epigenome feature extraction

Promoter DNA sequences were extracted around the TSS (−1000 nt to +50 nt) on the sense strand. CpG number was by CG dinucleotide, TATA number was by the “TATA” pattern. For counting ChIP-seq signal density, TSS intervals (−500 bp to 500 bp) and Ensembl (GRCm38.79) gene body intervals were used, with the samples as below:

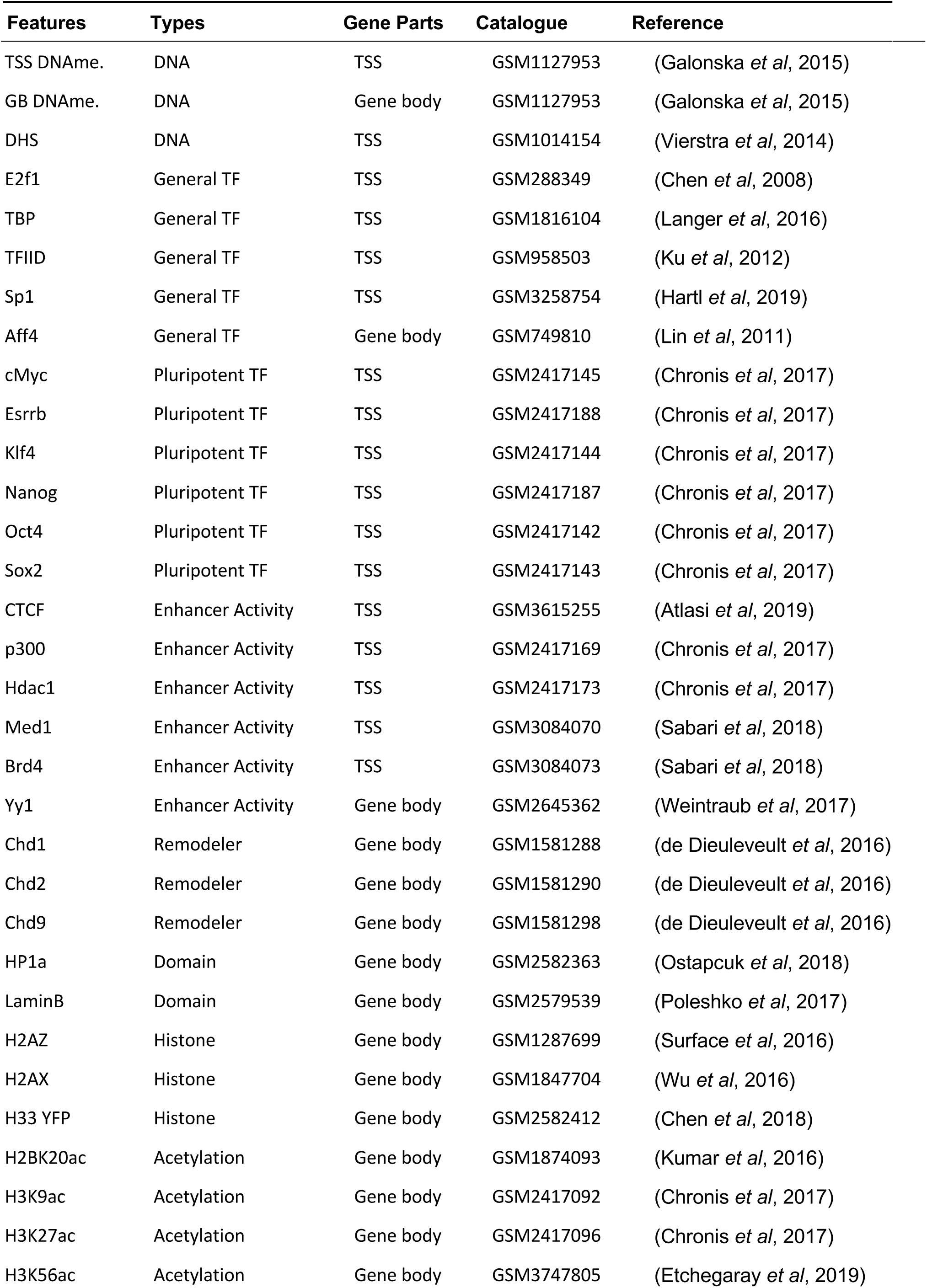
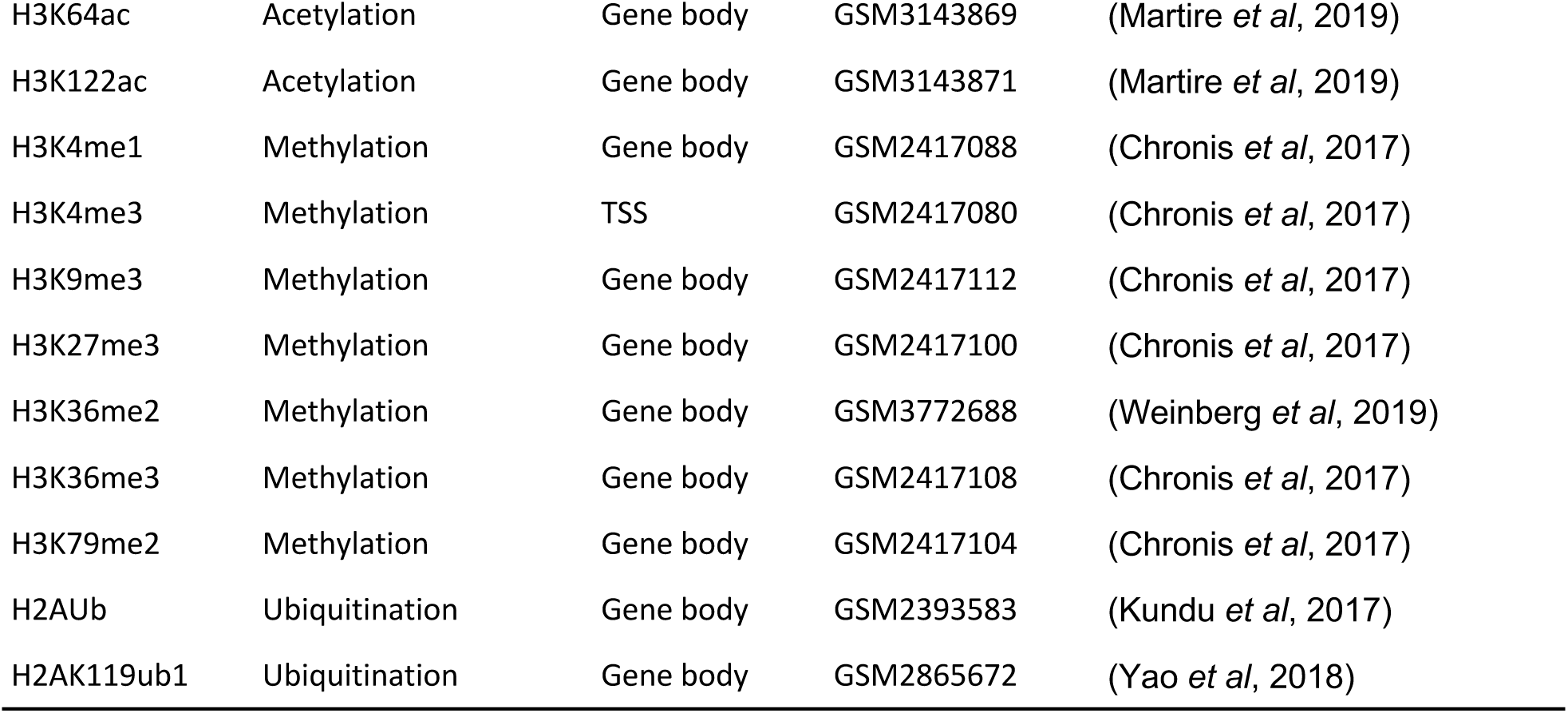

Log1p transformed ChIP densities were trimmed at 99.5% quantile to remove technical outliers, and standardized to normal distribution N(0, 1). Each feature explaining a particular response (e.g. read-through distance, Fig 5F) was decomposed for *R*^2^, by a multivariate linear regression through the origin.

### Pol II pausing index

Pol II TSS pausing intervals were generated from the closest 5’ capped RNA peaks (Start-seq (Dorighi *et al*, 2017), GSM2586572). Start-seq reads were aligned by STAR 2.7.3a and called peaks by HOMER 4.11 (Heinz et al.) with the following setting: findPeaks-style groseq-size 20 -fragLength 20 -inputFragLength 40 -tssSize 5 -minBodySize 30 -pseudoCount 1. Pausing intervals of the active gene were assigned by the closest non-redundant capped RNA intervals, which were used for TSS peak density calculation. Pol II S5p MINUTE-ChIP bigwig coverages, processed as described before (Kumar & Elsässer, 2019), were subjected to gene body density extraction from (TSS+500 to TSS+1500 bp) interval. The resulting ratios were used as the pausing index (Fig 3F).

### Transcription elongation velocity estimation

The time course Cdk9 inhibition GRO-seq samples (Jonkers *et al*, 2014) were annotated by TU filter to capture ongoing transcription events. The current travel distance from gene TSS were retrieved by the nearest TU fragment annotation, and were subjected to a linear regression model in response to Cdk9 inhibition times where the slope coefficient represented reversed elongation velocity and the intersection term adjusted for response time delay. The resulting elongation velocities for 3703 genes were used as the “measured elongation velocity” for cross-validation. Elongation velocities v can also be estimated from the ratio of the number of polymerases released into elongation, as measured by TT-seq, over the Pol II occupancy (Gressel *et al*, 2019; Caizzi *et al*, 2021). Thus, to derive “estimated elongation velocity” from our multi-omics data we combined TT-seq LRNA coverage with Pol II S5p MINUTE ChIP-seq coverage as follows:

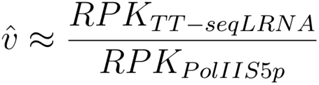

This approximation allows relative comparison between different conditions as long as the numerator and denominator terms, TT-Seq and Pol II S5p signals are quantitative. To this end, TT-Seq normalization by external spike-ins is performed as described above, and MINUTE-ChIP quantitative scaling is carried out as described (Kumar & Elsässer, 2019).

### Calling termination sites

For the 15 kb genome interval extending from the last exons of each gene, TT-seq LRNA reads coverage was piled up with the first paired read, log transformed and binned with 75 bp interval. To capture the significant decline of RNA synthesis after transcription passing the ultimate termination site, we slided the position that gave the maximum contrast of labeled RNA densities before and after that site. In practice, TT-seq nascent RNA read coverage in the termination window was scaled (mean = 0) and their cumulative sums from the beginning to the end of the termination window were calculated. The maximum position was defined as the ultimate termination site (Fig 5A).

### Non-linear model prediction of read-through distance

We used a tree-based gradient boosting model (gbm, R package) to evaluate the non-linear response of read-through distance with 41 chromatin features. The model training control was 10-fold cross validation, and with the tuned depth 40 and 1000 trees under 0.1 shrinkage. Test set was split by the ratio 0.2 of total cases (n=8348). The trained model of predicting numerical read-through distance was applied to the test set, and compared to the actual read-through lengths (Fig EV5D). Feature importance was extracted from this model as complement to the R-squared linear explanation (Fig 5C). We evaluated gbm performance with binned read-through groups (3, 5, 8, 15 kb) and ROC curves were then plotted to evaluate the model performance (Fig EV5E).

## Data availability

Primary and processed data generated for this study has been submitted to the Gene Expression Omnibus under GSE168378 (TT-seq) and GSE126252 (MINUTE-ChIP).

## Code availability

Transcription unit annotation R shiny app local workflow can be found on GitHub https://github.com/shaorray/TU_filter. Data analysis steps are also available on Github: https://github.com/shaorray/TT-seq_mESC_pluripotency.

## ACKNOWLEDGEMENTS

Bioinformatics analyses were performed on resources provided by the Swedish National Infrastructure for Computing (SNIC) at Uppmax server (projects SNIC 2020/15-9, SNIC 2020/6-3, uppstore2018208, SNIC 2018/3-669, sllstore2017057, SNIC 2017/1-508). We thank members of the Elsässer lab for comments and help with experiments and analysis. S.J.E acknowledges funding by the Karolinska Institutet SFO for Molecular Biosciences, Vetenskapsrådet (2015-04815, 2020-04313), H2020 ERC Starting Grant (715024 RAPID), Åke Wibergs Stiftelse (M15-0275), Cancerfonden (2015/430). R.S. acknowledges funding from the Chinese Scholarship Council.

## AUTHOR CONTRIBUTIONS

R.S. and S.J.E. conceived study. R.S. performed TT-seq and B.K. performed MINUTE-ChIP experiments. R.S. analyzed all data. K.F., M.L. and P.C advised on TT-seq. R.S. generated figures. R.S. and S.J.E. wrote the manuscript with input from all authors.

---

**Figure EV1.**
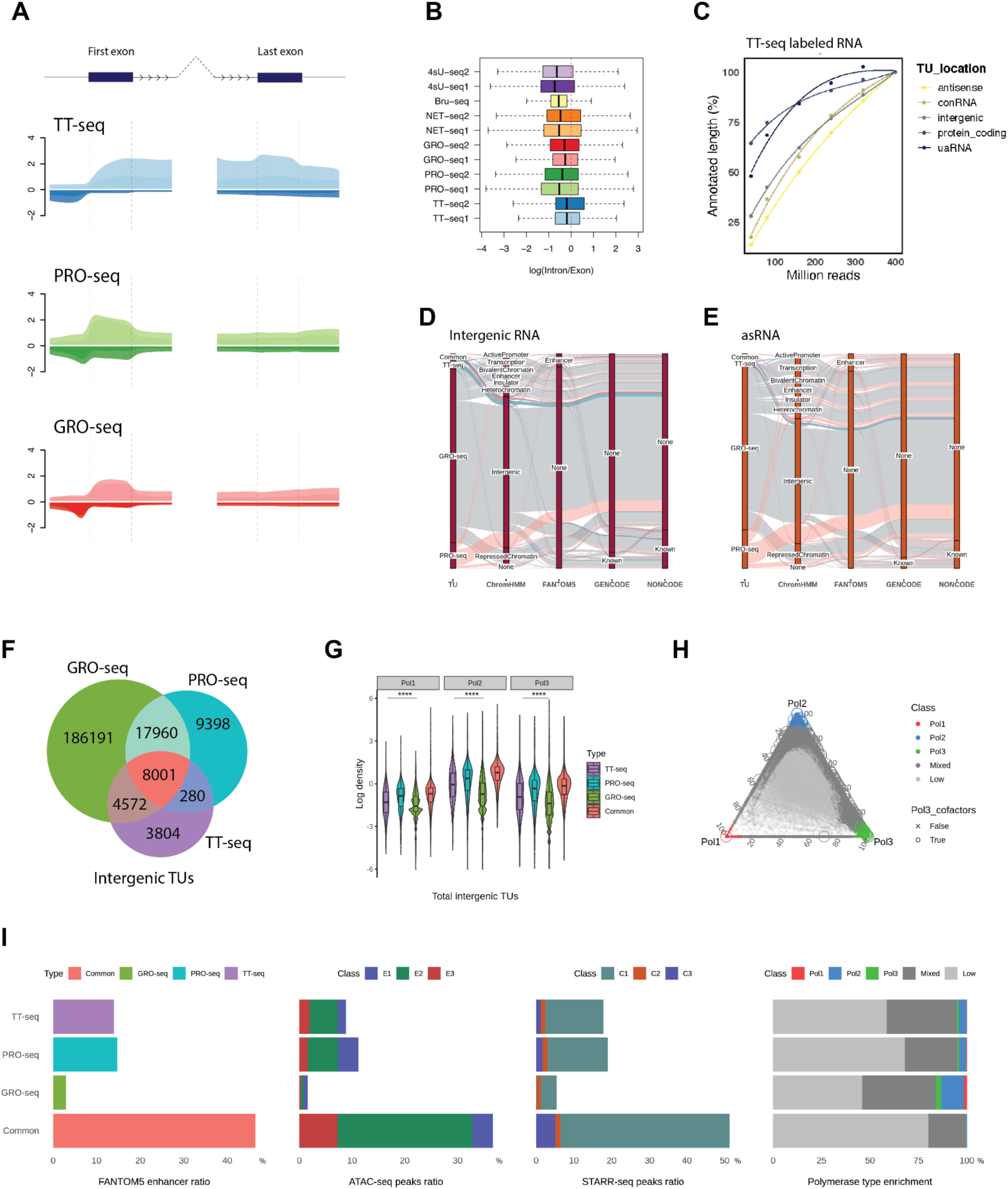
Nascent RNA-seq method comparison. A. Read coverages on the first and the last exons with 2 kb flanking regions and the adjacent introns (arrows). Exon and intron coverages were resized to the same dimension and plotted by log mean. B. The ratios of intron versus exon nascent RNA reads density on Ensembl protein-coding genes across different nascent RNA sequencing methods. C. TU annotation relative total length recovery test by TT-seq bam file random subsetting under (0.1, 0.2, 0.4, 0.6, 0.8, 1) for different TU types. D-E. Comparison of intergenic and cis-antisense RNAs from different nascent RNA-seq methods matching with public references (GENCODE(Frankish *et al*, 2021), FANTOM5 enchancer(Andersson *et al*, 2014), NONCODE(Zhao *et al*, 2016)). F. Venn diagram of GRO-seq, PRO-seq and TT-seq intergenic RNAs. G. RNA Pol I, II and III occupancy (Jiang *et al*, 2020) on the total intergenic TUs. Student’s t-test was performed with the common TUs against the method-specific TUs (****p < 2e-16). H. A ternary plot of Pol I, II and III enrichment on the GRO-seq, PRO-seq and TT-seq combined intergenic TU annotations. Pol III co-factors binding sites (Carrière *et al*, 2012) were pinpointed for Pol III class assignment cross-validation. I. Common overlapped and method specific intergenic RNAs proportions with FANTOM5 enhancers (Andersson *et al*, 2014), ATAC-seq peaks (Atlasi *et al*, 2019), STARR-seq peaks (Peng *et al*, 2020) and Pol I-III occupancy in H.

**Figure EV 2.**
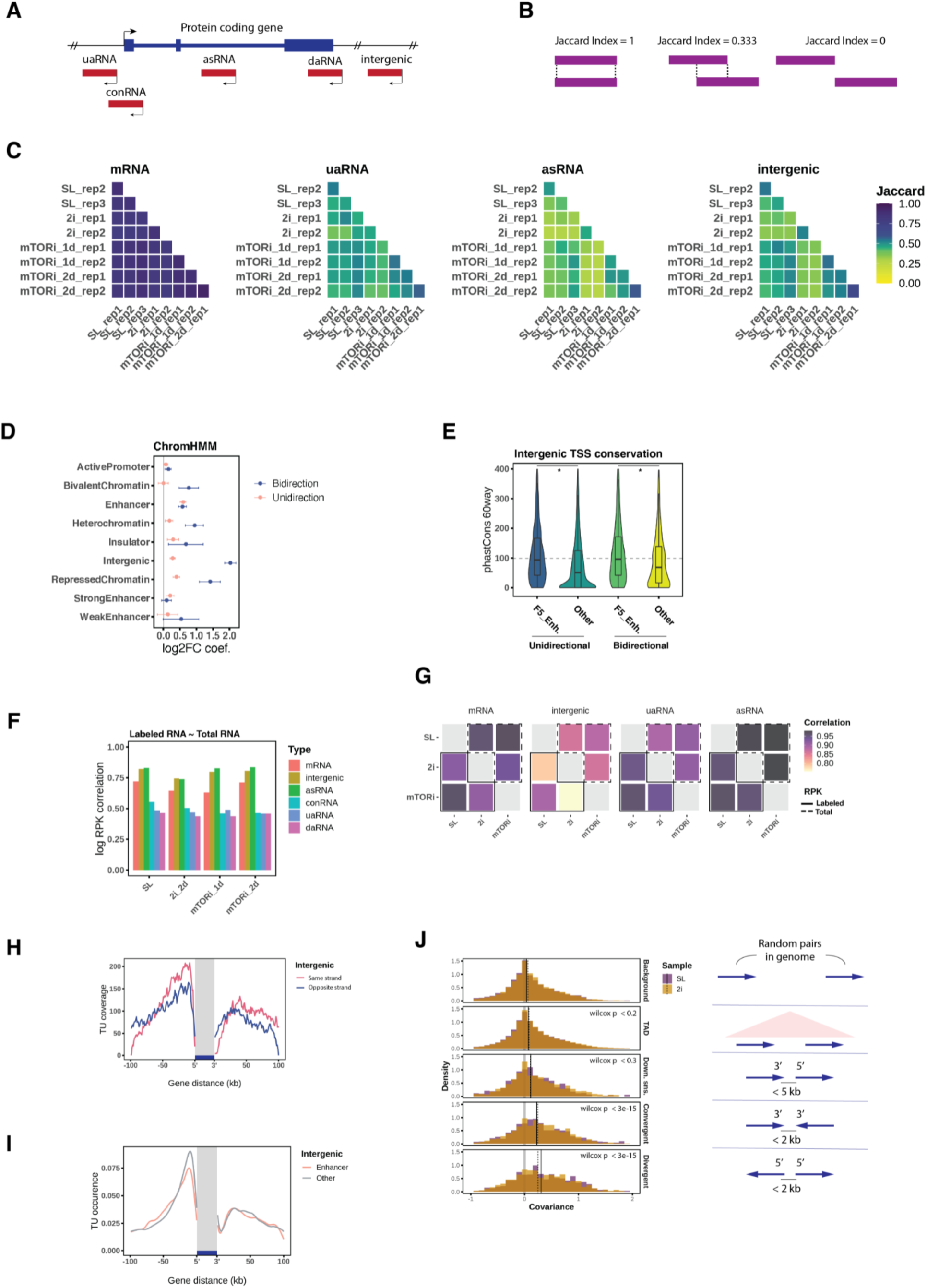
TU annotation and transcription variation. A. Gene associated ncRNA types were named by their TSS locations relative to the gene TSS: upstream antisense RNA (−1 kb, 0), convergent RNA (0, +1 kb), cis-antisense RNA (within gene body), downstream antisense RNA (TTS, TTS + 1 kb) and intergenic RNA. B. Illustration of Jaccard index of annotated TUs comparison. C. Jaccard indexes of mRNA, uaRNA, asRNA and intergenic RNA intervals similarity in pairwise comparison between each sample and replicate. D. The coefficients of ChromHMM states are in response to the internal normalised intergenic RNA log2FC differential transcription in 2i 2 days transition. Two separate logistic regression models were trained for the unidirectional and bidirectional intergenic TUs. Each states’ coefficients were plotted with the respective logistic regression confidence intervals. E. Intergenic TU promoter (−500 bp, 200 bp) evolution conservation scores from phastCons 60way (Siepel *et al*, 2005) with the same groups in Fig 2C (*Student’s t-test p < 2.2e-16). F. Pearson’s correlation between labeled RNA and total RNA by each TU location and pluripotent state with merged replicates and log RPK on the combined TUs, the same for G. G. Pearson’s correlation between the pluripotent states by labeled and total RNA log RPK. H. Intergenic TU intervals stack coverage in the gene ±100 kb neighborhood by relative strandedness. I. Intergenic TU TSS occurrence by enhancer (n = 3950) and other (n = 4081) states in genes ±100 kb neighborhood (n = 11684). J. Single-cell gene expression (Buettner *et al*, 2015) covariance distribution by the gene pairs of random background, within topological associated domain (TAD), consecutive downstream sense, 3’ ends convergence and promoter divergence. Wilcoxon test was performed against the background covariance. Median covariances of each gene positioning type were indicated for both SL and 2i states.

**Figure EV 3.**
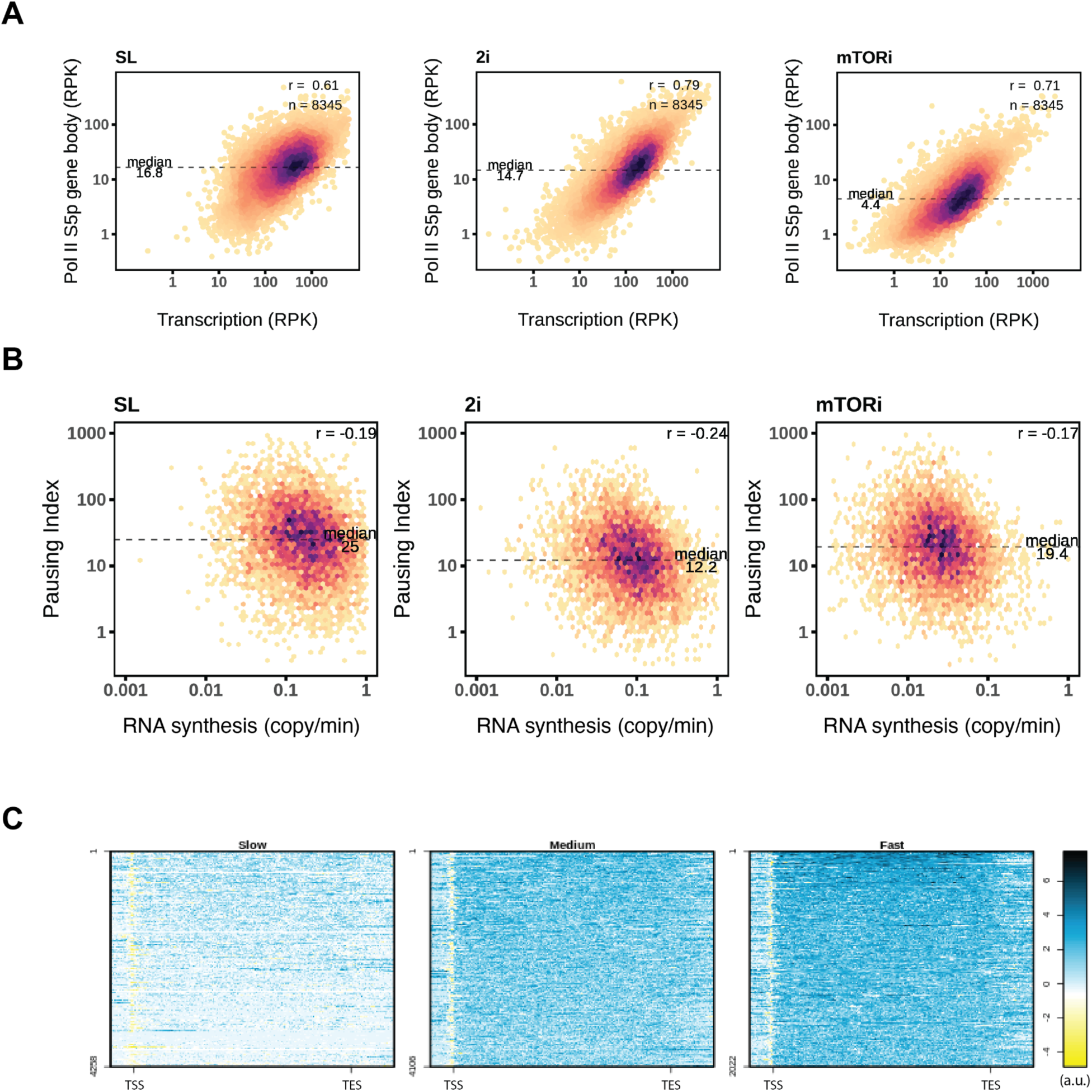
Pluripotent states transcription profiles. A. Scatter plots of Pol II S5p gene body occupancy and TT-seq nascent RNA reads density. Pearson’s correlation was performed on the log scale, the same as in B. B. Pausing indexes by Pol II S5p compared to RNA synthesis rates for each pluripotent condition. The medians of pausing indexes and Pearson’s correlation coefficients were indicated. C. K-means grouped estimated elongation velocity gene coverages were plotted in log scale. Upstream 2 kb and downstream 4 kb were extended from Ensembl protein-coding gene intervals.

**Figure EV4.**
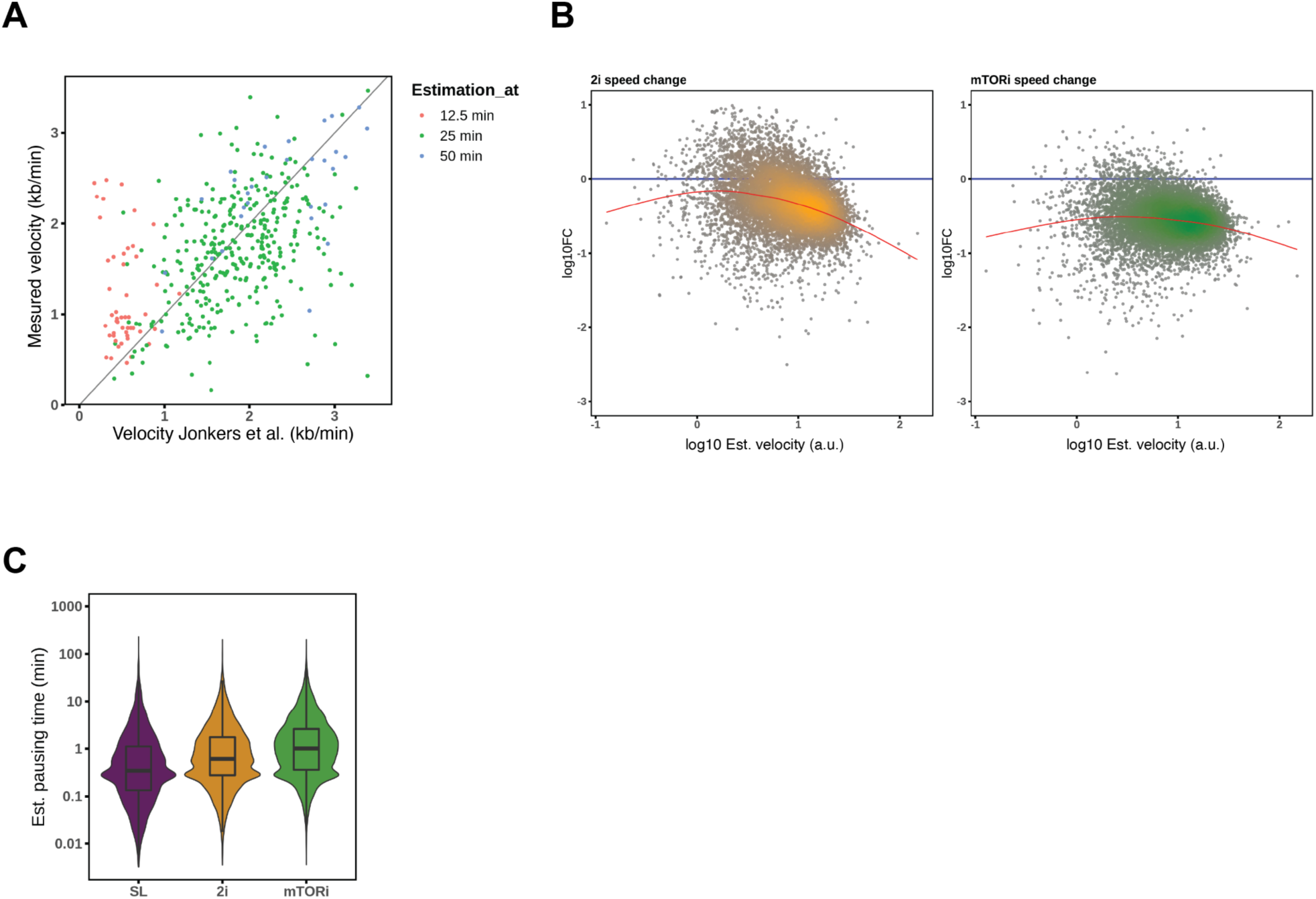
Pausing index and elongation velocity comparison. A. Comparison of the provided elongation velocity at the specified time points (Jonkers *et al*, 2014) and recalculated multi-time-point velocity estimates by linear regression (Materials and Methods). B. MA-plots of elongation velocity changes in 2i and mTORi conditions against estimated velocity in SL condition. Local regression lines were appended to illustrate the trend of changes. C. Estimation of pausing time in STAR-seq TSS intervals by the pluripotent states. Transcription velocity were scaled according to the external GRO-seq measured velocity.

**Figure EV5.**
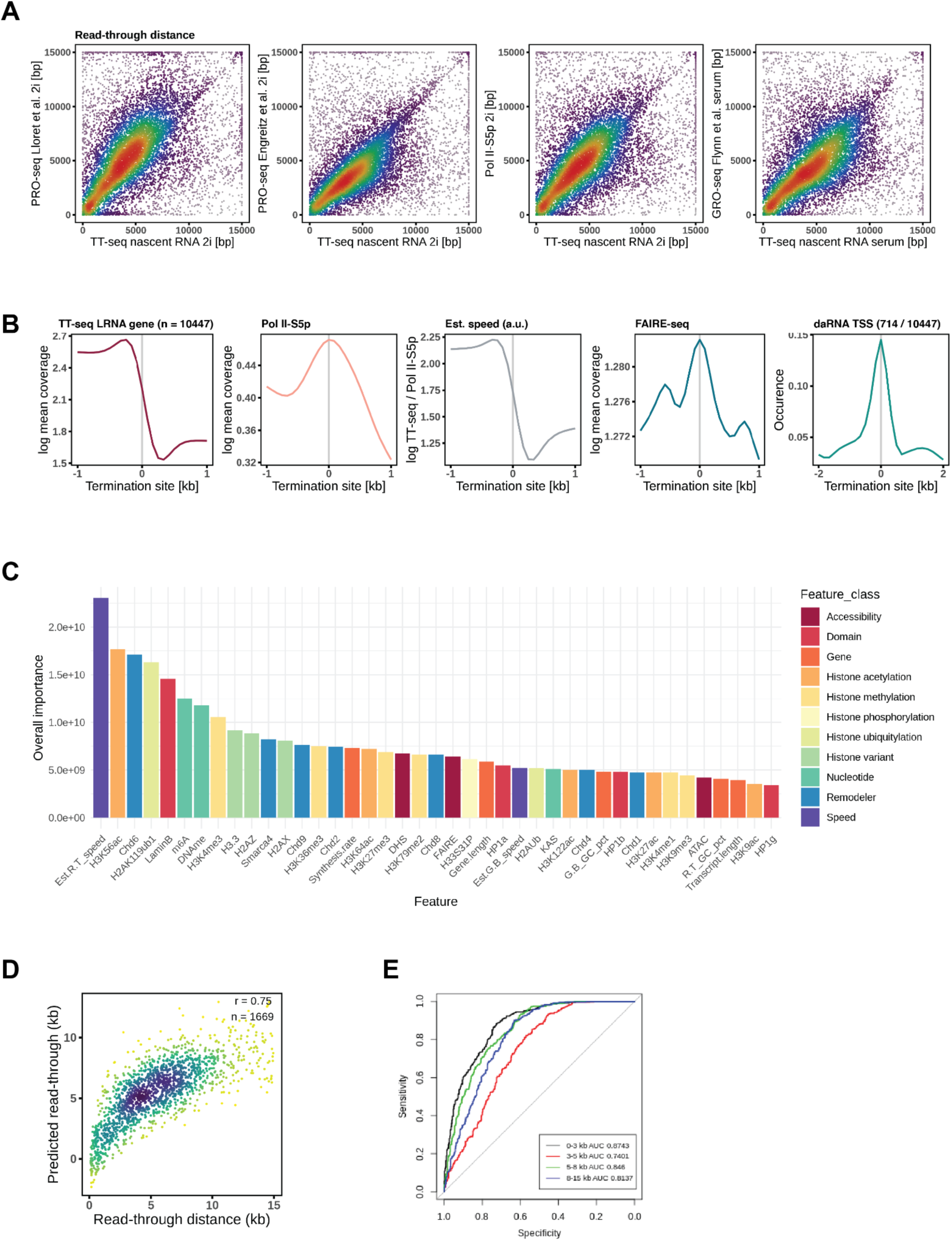
Transcription termination sites comparison and explanation. A. Read-through distances called by TT-seq, PRO-seq and Pol II S5p coverages in the termination window were compared by scatter plots. B. Average coverage around ±1 kb termination sites of TT-seq nascent RNA, Pol II S5p, estimated elongation velocity, FAIRE-seq and daRNA TSS occurrence with 10447 protein-coding genes. C. Gradient boosting machine (gbm) model’s feature importance of predicting read-through distance. 41 genomic features were used, the same as below. D. Comparison of the actual read-through distances and the predicted distances on the hold-out test set by the gbm model. E. A receiver operating characteristic curve (ROC) of showing read-through distance groups prediction performance with features as described above.

**Figure EV6.**
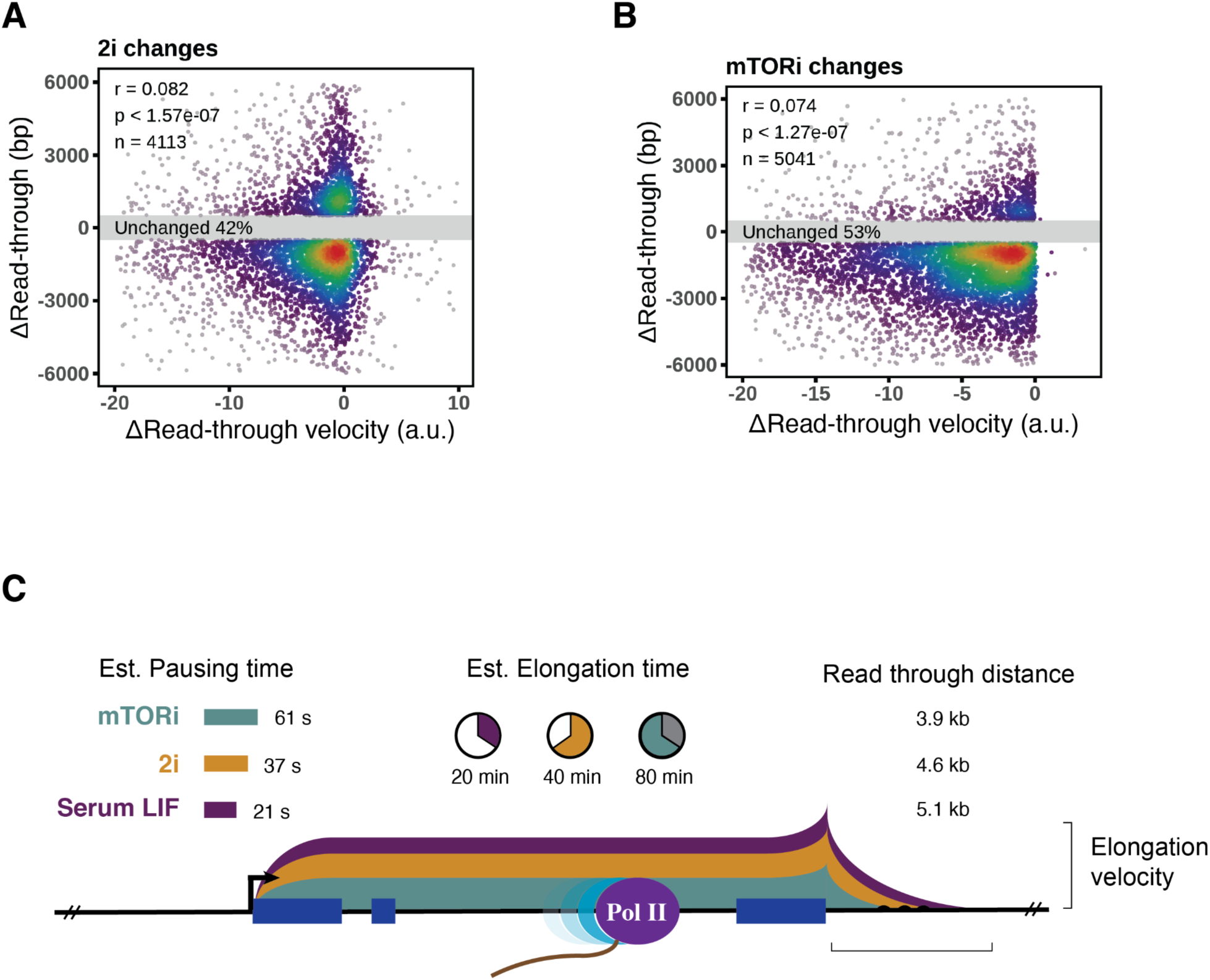
Elongation velocity and read-through distance association. A-B. Scatter plots of elongation velocity changes and read-through distance changes of 2i and mTORi transitions. C. Schematic of RNA transcription dynamics in mESC pluripotent states. Elongation velocity was scaled according to GRO-seq measurements, read through distances were from TT-seq termination window coverage.

